# Imagined Speech Reconstruction with 3D Neural Metabolism and Large Language Model Integration

**DOI:** 10.1101/2025.07.30.667805

**Authors:** Gowrish Rajagopal, Aniket Chaudhari, John LaRocco, Jinyi Xue, Eric Zachariah, Shwetan Varanasi, Douglas Rothman

## Abstract

Cognitive linguistics posits that language underpins human thought, and this principle has influenced the study and development of large language models (LLMs). In particular, several studies have investigated the metabolic costs of sentence formation using neuroimaging techniques such as positron emission tomography, functional magnetic resonance imaging, electroencephalography (EEG), and imagined speech reconstruction (ISR). In this study, EEG data corresponding to imagined English-language speech phonemes were used for ISR, in combination with an LLM trained on an abridged autobiography. The LLM-generated text responses guided the synthesis of EEG data from relevant phonemes, which were then used to estimate corresponding metabolic activity, and the changes in simulated neurometabolic and electrical parameters were visually represented. Notably, introducing pseudorandom variance significantly (*p* < 0.001) enhanced the model’s ability to reflect biological variability. Future directions include expanding the ISR system with lightweight or locally run LLMs, incorporating training data from larger and more diverse populations, and utilizing truly random variability sources. Further optimization for broader hardware compatibility and implementation—such as neural phantoms, emotional context integration, or human-computer interaction platforms—offer promising pathways for advancement. Overall, this work establishes a foundation for the next generation of biologically inspired, modular, and adaptable ISR systems for both research and practical applications.

## Introduction

### Overview

Cognitive linguistics asserts that language is the basis of thought, inspiring the design of various large language models (LLMs). LLMs, however, require significant amounts of energy and computational resources to run, whereas the human brain performs far more computations while using less energy. Previous studies have explored the metabolic costs of sentence formation using neuroimaging techniques such as positron emission tomography (PET), electroencephalography (EEG), and functional magnetic resonance imaging (fMRI). The use of imagined speech reconstruction (ISR) is one noted method (Tang et al., 2024). Rothman et al. (2021) experimentally validated a model of human brain metabolism, effectively estimating neural metabolism and converting between neuroimaging modalities. As phonemes are the basic components of language, their production lies at the end of speech formation, and given that EEG data from the imagined speech of English-language phonemes is publicly available, it can easily be encoded into a text string. Using a ‘fractal’ brain model, software was developed to convert any text string into a scalable resolution, 3D model of neurometabolic activity based on ISR.

### Background

#### Cognitive Linguistics

Consciousness itself cannot be reliably quantified, but discrete patterns of brain activity can (Cogitate Consortium et al., 2025). Investigating the relationship between language and the brain, particularly the speech centers in general cognition, reveals the intricate neural landscape involved in language processing. The left hemisphere’s predominance in language function is well known, especially within certain regions such as Broca’s area, Wernicke’s area, and the angular gyrus. Evidence indicates that these regions are involved not only in speech production but also in semantic processing and comprehension, illustrating a network architecture that is critical to linguistic capabilities (Price, 2012; Clercq et al., 2024; Imaezue, 2017).

Broca’s area, located in the left frontal lobe, is primarily associated with speech production and syntax, interacting continuously with the temporal and parietal lobes. The temporo-parietal junction is crucial for auditory language perception and production, with increased activation observed during speech-related tasks compared with non-linguistic stimuli (Price, 2012; Krishnan & Gandour, 2009). Notably, a meta-analysis revealed that both first and second languages recruit similar neural pathways, suggesting that proficiency impacts processing but does not fundamentally alter the brain regions involved (Sebastian et al., 2011). This demonstrates the robustness of the neural representations associated with linguistic tasks.

Moreover, functional brain connectivity has emerged as a key research area, linking disparate regions across the left hemisphere during language tasks. For instance, studies utilizing fMRI have shown synchronous activation patterns between Broca’s and Wernicke’s areas, thus highlighting a functionally interconnected network essential for fluency and comprehension (Tripathi, 2024; Fedorenko et al., 2011). The interaction between these areas supports the processing of phonological, syntactic, and semantic information, all of which are necessary for effective communication (Fedorenko & Thompson-Schill, 2014; Friederici, 2004).

Recent advancements in the understanding of how bilingualism affects brain morphology indicate that language processing is significantly shaped by an individual’s language experience, resulting in adaptive neural structures (Burgaleta et al., 2016; Guan, 2024). These structural changes not only enhance linguistic capabilities but also facilitate cognitive flexibility, underscoring the intertwined nature of language use and general cognitive functions.

Furthermore, research has shown how language processing engages domain-general cognitive resources, suggesting that some brain areas function in both linguistic and non-linguistic tasks (Fedorenko et al., 2011; Heuven et al., 2008). This overlap indicates that while certain regions are specialized for language, they also partake in broader cognitive operations, thus blurring the lines between language-specific and general cognitive functions.

In summary, language generation involves a sophisticated, highly interconnected neural framework in the brain that governs both speech production and comprehension. This architecture not only facilitates language-specific processes but also contributes to a spectrum of cognitive functions, illustrating the profound complexity of human language processing, as mediated by brain structures.

#### Brain Metabolism Measurements

Precisely measuring neural energy expenditure is a complex, highly inconsistent process (Saveliev et al., 2023). Prior work on scalable simulations of brain metabolic molecule concentrations has provided significant insights, utilizing advanced imaging and analysis techniques to explore metabolic variations across different brain regions. A comprehensive metabolome atlas was created by Ding et al. (2021), establishing a foundation for understanding the metabolic differences present in specific mouse brain regions, particularly in relation to aging. Their findings revealed that these metabolic variations are closely associated with functional and molecular changes and can serve as a reference for health assessments in various states, including disease.

Building upon this foundational work, several studies have employed imaging techniques to correlate metabolic activity with specific brain pathologies and neurophysiological states. For instance, Jacob et al. (2019) conducted a study using fluorodeoxyglucose to demonstrate significant decreases in metabolic activity in specific brain regions affected by binge drinking, effectively illustrating the application of regional imaging in clarifying the functional consequences of external influences on metabolism. Similarly, Kleinridders et al. (2018) utilized imaging mass spectrometry to elucidate regional differences in glucose metabolism across the brain, finding that metabolic responses in specific areas like the thalamus and cortex were markedly distinct, thereby emphasizing the necessity for region-specific considerations in metabolic research.

Other research has utilized targeted metabolomics and imaging techniques to explore metabolic processes in relation to health states. For example, Zhou et al. (2022) examined oxygen consumption across various brain regions, revealing that changes in neural activity directly correlated with altered metabolic demands and oxygen levels, thus fostering an understanding of how metabolism varies dynamically with activity levels. Furthermore, Képes et al. (2021) observed the effects of metabolic syndrome on brain glucose metabolism, linking regional metabolic activity with broader health implications.

Technological advancements have also enabled high-resolution methodologies for mapping brain metabolism. Pang et al. (2020) provided a detailed visualization of metabolite distributions across the mouse brain, significantly enhancing our capacity to interpret metabolic data based on brain architecture. This represents a critical advancement in integrating spatial resolution with metabolic profiling in simulations.

Additionally, simulation frameworks like BrainScale have been developed to address the complexities associated with large-scale spiking neural networks, highlighting the importance of robust computational methodologies for advancing neuroscientific research (Wang et al., 2024). These frameworks facilitate the incorporation of metabolic models, allowing researchers to simulate brain metabolism in a scalable and efficient manner.

In summation, the confluence of metabolomics, advanced imaging, and computational modeling has yielded a robust landscape for the examination of brain metabolic processes, showcasing diverse regional variations and their implications for understanding brain function in health and disease. Continued exploration in this field is essential for unraveling the complexities of brain metabolism and its role in cognitive and neurodegenerative disorders.

#### EEG as a Metabolic Marker

The validation of mathematical models to predict metabolic concentrations in the brain from EEG data is an evolving field that integrates insights from neuroscience, machine learning, and clinical practices. EEG is a simple, accurate, and non-invasive tool for assessing brain metabolic states and predicting associated metabolic profiles.

Numerous studies suggest that EEG can be effectively linked to metabolic processes in the brain. For instance, Westover et al. (2015) proposed a method for estimating brain metabolic state based on EEG burst suppression, demonstrating how electrophysiological features correlate with cerebral metabolic rates. This indicates that EEG data can be used to determine metabolic conditions. Similarly, Schamberg et al. (2020) developed a model that represents the transition dynamics among burst and suppression epochs obtained by EEG, which is influenced by the metabolic level and quantified based on ATP production.

Moreover, Crouzet et al. (2016) argued that understanding the discrepancy between EEG signals and metabolic deficiency post-cardiac arrest leads to improved predictive modeling regarding EEG recovery and neural function, emphasizing the importance of integrating EEG reactivity and metabolic assessments for better outcomes in patients with neurological impairments. Understanding how EEG features can signal metabolic deficiencies allows for enhanced prognostication strategies and therapeutic decisions in critical care settings (Müller et al., 2020; Lee et al., 2019).

Machine learning approaches further augment the predictive capabilities of EEG data for metabolic states. The Auto-Neo-EEG model introduced by Dong et al. (2021) employs machine learning to assist in clinical EEG report generation, thereby reinforcing the predictive precision achievable with advanced computational techniques. Likewise, McConnell et al. (2024) demonstrated that digital biomarkers derived from EEG data can be leveraged to predict various health metrics.

In addition, integrating EEG with other modalities, such as functional near-infrared spectroscopy (fNIRS), can enrich models for predicting brain metabolic states and vascular coupling (Pinti et al., 2021). This interdisciplinary approach highlights the importance of using a multimodal perspective when modeling brain activity and metabolic functions.

Ultimately, ongoing research in EEG methodologies, combined with robust mathematical and computational models, paves the way for reliable predictions of metabolic concentrations in the brain. Such developments not only enhance clinical monitoring capabilities but also hold transformative potential for understanding brain health across various physiological and pathological states.

#### Brain Metabolism Models

Scalable simulations of brain metabolic molecule concentrations have gained prominence in integrating neuroimaging techniques like EEG, fNIRS, and fMRI, providing a more comprehensive understanding of brain metabolism in health and disease. In particular, PET imaging, especially using 18F-fluorodeoxyglucose, has been pivotal in mapping regional glucose metabolism across various conditions. In the investigation of brain metabolic networks, alterations in glucose uptake can indicate different pathological states, such as those observed in Parkinson’s disease (Tan et al., 2018; Li et al., 2020; Matthews et al., 2018).

The integration of functional neuroimaging has shown that metabolic abnormalities in regions like the putamen and globus pallidus correlate with clinical symptoms in Parkinson’s patients, demonstrating how metabolic assessments can guide therapeutic strategies (Tan et al., 2018; Li et al., 2020). Furthermore, machine learning has been used to develop predictive models based on metabolic networks, linking brain metabolism to functional outcomes in cancer patients, which suggests that brain metabolism is influenced by systemic health conditions (Yu et al., 2023).

Experimental validation reinforces these insights. For example, magnetic resonance spectroscopy enables the non-invasive monitoring of metabolites such as N-acetylaspartic acid, GABA, and glutamate, all of which have been linked to specific neurological functions and dysregulations in various diseases (Mao et al., 2024; Hyder & Rothman, 2017). Neurometabolic coupling, which can be examined using simultaneous imaging techniques, such as spatial frequency domain imaging, is critical for evaluating brain function, particularly during intervention studies (Wilson et al., 2019).

EEG and fNIRS have additionally been utilized to detect changes in cerebral glucose metabolism during cognitive tasks, reinforcing the link between neural activity and metabolic processes (Ghijsen et al., 2018; Crouzet et al., 2016). Advancements in imaging technology allow for real-time analysis of brain metabolism, contributing to the comprehension of conditions like Alzheimer’s disease, where altered glucose metabolism precedes clinical symptoms (Camandola & Mattson, 2017; Goyal et al., 2023). Novel imaging methodologies, such as dynamic deuterium magnetic resonance spectroscopic imaging, facilitate the concurrent measurement of multiple metabolic rates in vivo, offering a detailed view of brain energy dynamics (Tolomeo et al., 2018).

The combination of imaging modalities such as PET and MRI has also been transformative, yielding comprehensive data on metabolic activity and structural integrity (Li, 2025; Radder et al., 2024). This fusion allows for improved characterization of brain pathologies and enhances the precision of targeted therapies by integrating functional, structural, and metabolic information within a single framework. Similarly, integration with advanced visual display software is expected to increase our capabilities in simulating and interpreting brain metabolic processes.

Overall, advances in scalable simulations and their integration with neuroimaging techniques provide profound insights into the realms of brain metabolism, correlating various molecular pathways with observable physiological phenomena in both healthy and diseased states. Regarding the present study, simulating known metabolic pathways can enable direct estimation of the neural energy and processing costs, especially during sentence formation. Such a tool has direct implications for neurolinguistics and ISR, although simulating the active variability in biology has proven difficult in prior work (Pliatsikas et al., 2021; Guan, 2024; Tang et al., 2024).

## Methods

### Pipeline Design

Neurolinguistic metabolistic simulations were performed in this study, producing metabolic simulations from text string inputs. The pipeline contains custom-built modules that integrate language modeling, phoneme-based EEG synthesis, computational metabolic modeling, and 3D brain renderings to offer a biologically based presentation of how cognition and physiology dynamically respond to speech-based stimuli. The system was trained with publicly available datasets and documents, as a template for further development.

The pipeline is summarized in Figure 1. The overall process begins with a personalized GPT-4 conversational agent that simulates the style of an abridged autobiography (OpenAI, 2025). The responses are then phonetically decomposed and mapped to an EEG dataset of phoneme-label recordings, acquired from a public dataset (LaRocco et al., 2023). The synthesized EEG sequence is then processed through both neuroelectrical (MNE-Python-based source localization) and neurochemical (gamma-band to CMRO_2_ translation) modeling (Pliatsikas et al., 2021). A set of user-selected neurovascular variables (including cerebral blood flow (CBF), pH, and partial pressures) is computed and projected onto a cortical surface via interpolation, weighted by electrode proximity. Animations are produced to visualize electrical activation metabolic changes in the brain throughout the GPT-generated response, all while providing a dynamic user experience through the graphical user interface (GUI).

**Figure 1:**
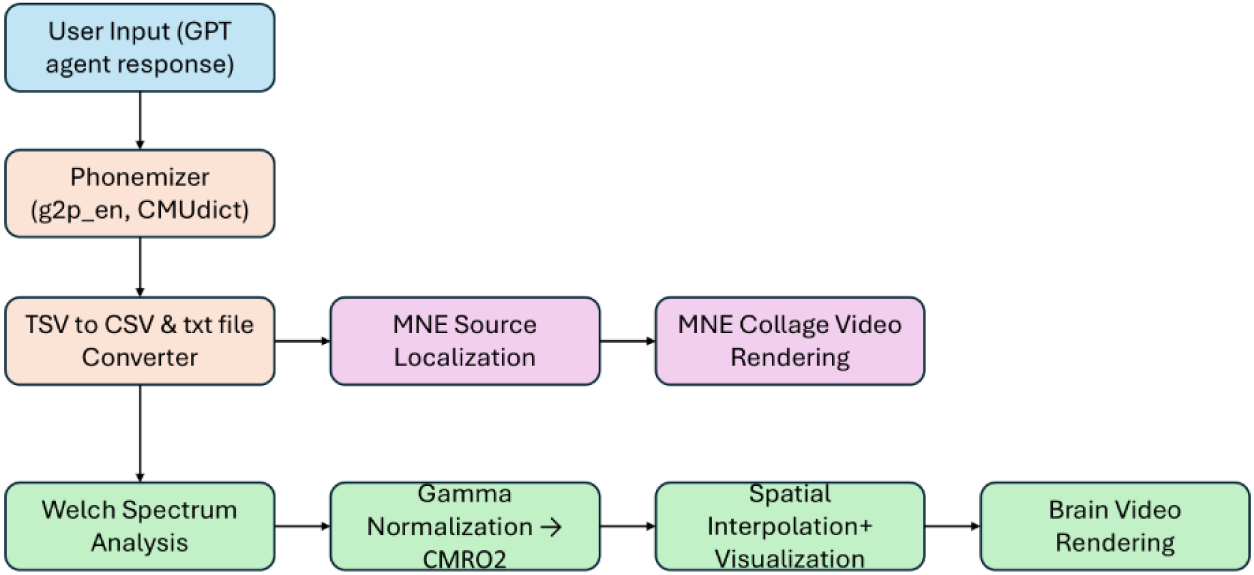
Processing pipeline graphical summary.

### Conversational Language Generation

To begin the process of neuro-visualization modeling, a conversational agent based on GPT-4 was created using the LangChain framework (LangChain, San Francisco, California, United States). Multiple components were incorporated to enhance the conversational flow between the agent and the user, including the ability to return and retain context from previous messages with RunnableWithMessageHistory. These additions led to richer and more coherent responses in the style of the provided text.

User inputs were passed through a memory-aware retrieval chain backed by a Facebook AI Similarity Search vector database (Meta, Menlo Park, California, United States). The retrieval system populated the vector store with embeddings from OpenAI’s text-embedding-large model applied to pre-authored texts (OpenAI, 2025). Retrieved information found in the texts was fractioned and assessed through a combined_documents_chain to provide related context for modifying the agent’s output.

The GPT-4 model was guided by prompt engineering to mimic the thought process and linguistic style of the author’s introspection in the text. The prompt limited the agent’s responses to 20 words and instructed the agent to provide personalized messages in a curious and scientific tone. The overall system consisted of both a response generator and a question reformulator, allowing follow-up queries to be passed and interpreted as independent prompts without undesired characteristics. The final responses were formulated by the system and rendered on the GUI to be used as input for the phoneme-to-EEG phase of the pipeline.

### Phoneme Decomposition

The response text was first cleaned by removing extraneous punctuation and white space at the word level. Each word was then passed onto a hierarchical phoneme extraction system that conducted a dual-layer lookup process. Specifically, each word was searched for in the Carnegie Mellon University Pronouncing Dictionary (CMUdict, Carnegie Mellon University, Pittsburgh, Pennsylvania, United States), which contains a corpus of over 100,000 words and their corresponding phonemic transcriptions in ARPAbet format. To resolve cases where the word was not found due to capitalization or non-English origin, a neural grapheme-to-phoneme model (g2p_en, Anaconda, Inc., Austin, Texas, United States) was employed, generating inferred ARPAbet phoneme combinations based on raw text input.

The resulting phoneme sequences were then standardized. ARPAbet transcriptions tended to include stress markers (e.g., “AE1” or “AA2”) to portray emphasis on specific syllables, but these were not relevant for EEG waveform pairing. Accordingly, the markers were stripped to produce a stress-invariant phoneme list. This design choice prioritized signal consistency over stress variation, given the nature and limitations of the available EEG dataset. The complete set of standardized phonemes was displayed to the user, employing an EEG segment retrieval process to assemble a synthetic yet biologically based neural representation of the original GPT-generated prompt. Users could also access voice recordings of the generated sentence output and the individual phonemes, which were synthesized using pyttsx3 (Anaconda, Inc., Austin, Texas, United States).

### EEG Segment Assembly

Once the phonemes had been standardized, each element in the sequence was mapped to a corresponding segment from a phoneme-labeled EEG dataset collected from a participant under controlled laboratory conditions (LaRocco et al., 2023). The EEG recordings were acquired using a 16-channel configuration (Fp1-O2), with each phoneme corresponding to a 1-second segment sampled at 256 Hz. The recordings were used interchangeably to create temporally consistent phoneme-specific neural responses.

Each phoneme in the sequence was passed through the system, which retrieved the represented EEG file from a pre-indexed dataset using a numerical ID mapped to each ARPAbet unit (e.g., “B” → 7 → DLR_7_1.txt). The retrieved EEG file was subsequently parsed line-by-line to locate the first occurrence of the 0th index value, indicating that the first complete 256 samples aligned with the phoneme event. The first second of data (256 samples) was taken from the starting index and appended to an output buffer. A warning was logged if the required EEG file was missing or incomplete, and the corresponding phonemic segment was skipped in the final composed file.

To smoothen the discontinuity between words and maintain natural temporal structure, pseudorandomized “microgap” segments were inserted between words. The gaps were chosen by randomly sampling from a set of five additional EEG recordings within the same dataset and were randomly selected to be either 256 or 512 samples long, with an equal probability, introducing variability similar to natural pauses and spoken fluctuations seen in human speech. With this addition, EEG segments became less abrupt, and the dynamics of sentence-level brain activity were better preserved. EEG segments were concatenated into a continuous stream of voltage values across the aforementioned 16 channels, creating a synthetic EEG representation of the original GPT-generated text.

### Visualization in 3D

The main process involved the visualizer.py script, using the MNE Python library to process EEG data from a CSV file, perform source localization, and then visualize the results in the form of a 3D brain model (Gramfort et al., 2013). All the logic was encompassed in the class EEGVisualizer, taking the CSV file path as the input for the constructor. A class-object logic was chosen for this component because it could be utilized in any other Python file where only a CSV input was needed, making integration easier and faster. The class properties included the EEG channels, the sampling frequency at which the EEG scan was recorded, the MRI-based options, and the number of CPU cores needed to customize the computing power and processing time.

The visualize(self) method was implemented to read the CSV file, and the data was cleaned to obtain only the EEG’s numeric readings, which were converted to a transposed numpy array with the following configuration: (n_channels, n_samples). After the data was stored, a mne.info object was created to store necessary information related to the EEG data. The mne.info object was required for creating an MNE Raw object, the primary container for storing continuous raw EEG data. Then, the reference needed to calculate the EEG data was set to ‘average,’ and the signal was processed at each electrode and re-referenced by subtracting the average signal of those electrodes at each time point. This preprocessing step reduced noise in the data and improved the signal quality. Next, the MNE Raw object stored a standard montage that defined the 3D locations of the EEG electrodes on the scalp. Regarding the EEG headset, the 10-20 International System was used, as described in a previous study (Larocco et al., 2023).

With all the information for processing received, a source space was defined—a set of points within the brain volume where the electrical activity would be estimated. The density of the points was defined as ‘ico4,’ which represented a moderate number of points inside the brain volume. Then, a boundary element model was generated that defined the different tissues in the head and their electrical conductivity. All this information was fed to a forward solution that computed a lead-field matrix and modeled how EEG electrodes on the scalp would measure a current dipole at each point in the source space.

A covariance matrix was also computed based on the MNE Raw data to describe the statistical relationship between the electrode signals. This was essential for creating an inverse matrix later in the process. Specifically, a covariance matrix of [length of the CSV file × 16] was created, where the length of an input CSV file ranged from 500 to 40,000. Considering that this may take a long time to compute, the covariance computation was performed dynamically to choose which covariance method to use based on the length of the data.

The inverse matrix, which took the forward solution, the covariance matrix, and the MNE Raw object, was applied to the raw EEG data to estimate the activity in the source space. The inverse matrix’s data was normalized by their variance and applied to the raw data with a regularization parameter, thus creating a source time courses (STC) object that represented the estimated activity at each source point over time.

Using MNE, a brain model was generated with specific stylistic parameters, and then the STC object’s left and right hemisphere data were added to the model in segments. The data was cropped to one-hundredth of a second, and the rounded time of the total length of the EEG CSV file was divided by the sample frequency at which the EEG data was recorded. The output was an animated 3D brain model that showed the intensity of the electrical data at each time point inside the brain, based on the input EEG file.

### Visualizing Neurochemical Metabolism

The synthesized EEG file was sent into the first of two final phases of the pipeline, namely the production of an animated visualization of the neurochemical response across the cortical surface. The model simulated how the phoneme-derived electrical activity reflected metabolic dynamics. Brain activity is energetically demanding, and oxygen is a primary driver in energy production; therefore, electrical activity production serves as a means to estimate the activity and metabolic levels of different brain regions.

Similar to the previous study, the process began with the spectral decomposition of the EEG signal using Welch’s method found in the EEG_Implement_Welch file (LaRocco et al., 2023). The inputted EEG data was segmented into overlapping windows to calculate the power spectral densities. In this system, prioritization was placed on the gamma frequency band (30–100 Hz), which is known to correlate with metabolic demand. For each trial and electrode, gamma-band power was normalized either globally or per trial, depending on the user’s choice. These values were then mapped to estimates of the cerebral metabolic rate of oxygen consumption (CMRO_2_) through a linear transformation, where 2.1 μmol/g/min was considered the baseline metabolic rate, using the plot_normalized_gamma_across_channels module.

The estimated CMRO_2_ values were inputted into the calculate_neurovascular_variables function, based on a physiologically detailed model inspired by prior and related work on neurovascular coupling (Pliatsikas et al., 2021). Using standard values for hemoglobin affinity (P_50_ = 27 mmHg), buffering constants (BC = 26.0), and arterial pressures (pO_2_ = 100 mmHg, pCO_2_ = 40 mmHg), the following variables were calculated:

- Cerebral blood flow (CBF) was calculated from the relationship CBF = CMRO_2_ / (O_2_ concentration × OEF).
- Oxygen extraction fraction (OEF) was modulated using the coupling constant n = 2.28.
- pCO_2__V and pO_2__cap—the venous and capillary gas pressures of carbon dioxide (CO_2_) and oxygen (O_2_)— were calculated via mass transfer and hemoglobin binding models.
- pH_V—the venous pH—was estimated from lactate and CO_2_-related acid-base buffering.
- ΔHCO_2_ and Δlactate served as proxies for metabolic compensation and anaerobic flux.

Outputs were subsequently generated per electrode, per timestep, and per trial (if selected), creating a 4D matrix (variable × spatial × temporal × repetition) that helped simulate dynamic cortical biochemistry. The EEG_Plotting module was used to display the calculated data. The 16 physical electrode positions were placed using Latin hypercube sampling (n = user-inputted number of virtual nodes, often exceeding 1000) for dense surface populations (Loh, 1996). Red dots represented actual electrode placements, whereas blue dots indicated interpolated locations placed throughout the brain model to simulate the rest of the cortex. Each point was assigned a value using the DistanceFunc module, which computed the 4-nearest neighbor weighted averages based on Euclidean distances in the 3D hemispheric space.

In the end, users received a 3D animation in which the selected metabolic variable was rendered frame-by-frame across 32 time slices, forming a temporal progression of the chemical values. The chemical values were interpolated across the cortical space through electrode-weighted proximity for each frame. The surface was shown using a color scale with ranges determined based on biological markers observed in typical brain activity (e.g., CMRO_2_: 2.0–10.0, pH: 6.5–7.4, CBF: 20–90 ml/100 g/min). Each animation was saved as a .mp4 file and played through the GUI, allowing for the reselection of metabolic variables and a refresh of the phrase input created by the GPT agent.

### Statistical Tests

GPT-generated text was produced with and without pseudorandom space insertion from the full pipeline. Trials were subsequently processed to extract electrical and metabolic variables across 16 cortical channels per metric. Paired statistical analyses were conducted across all neurovascular measurements using parametric and nonparametric methods. Effect sizes and residuals were calculated to determine the magnitude and significance of the observed variations. Visual outputs included GUI renderings throughout the pipeline process, metabolic and electric overlay maps, and regression collages that portrayed the transformation of phoneme-converted EEG activity to modeled cortical metabolism and electrical activity. These visualizations represented a single trial, illustrating the detailed spatial resolution of the outputs.

The primary comparison was made between the GPT-generated texts with and without pseudorandom spacing across 3 trials. The values from each trial were averaged across the 16 cortical channels for each neurovascular metric (CBF, OEF, pH_V, pCO_2__V, pO_2__cap, CMRO_2_, ΔHCO_2_, and ΔLAC). Paired t-tests and Wilcoxon signed-rank tests were used to determine the significance of the spaces, and Cohen’s d was calculated to estimate the effect size. Residual means and standard deviations were computed to quantify the consistency of differences across the channels. Considering the higher potential variability, it was hypothesized that the differences with pseudorandom spacing would be significant.

## Results

### Overview

Three outcomes are presented herein. The first is the outcome of the statistical tests, the second is the presentation of the GUI, and the third is the output of the 3D brain plots.

### Neurovascular Variable Statistical Tests

Paired t-tests were performed across 3 trials for each neurovascular metric (CBF, OEF, pH_V, pCO_2__V, pO_2__cap, CMRO_2_, ΔHCO_2_, and ΔLAC).

#### Trial 1 Outcomes

The Trial 1 prompt and response are shown in Table 1.

**Table 1:** Trial 1 Prompt and Response.

|  |  |
| --- | --- |
| User Prompt: | What aspect of brain-computer interfaces (BCIs) do you find most compelling or exciting? |
| GPT-generated Sentence: | “The potential for BCIs to restore lost functions, like movement or speech, in individuals with disabilities. It’s a tangible bridge between technology and humanity’s resilience.” |

The statistical outcomes for Trial 1 are shown in Table 2. Differences were significant (*p* < 0.001).

**Table 2:** Trial 1 Statistical Outcomes.

| Metric | t_statistic | p_value_ttest | p_value_wilcoxon | cohens_d | residual_mean | residual_std | n_pairs |
| --- | --- | --- | --- | --- | --- | --- | --- |
| CBF | 4.602911283 | 0.000344985 | 0.000305176 | 1.150727821 | -5.00E-16 | 0.009894423 | 16 |
| OEF | -4.92115218 | 0.000184581 | 0.000305176 | -1.230288045 | -3.47E-18 | 7.71E-05 | 16 |
| ph_V | -4.92115218 | 0.000184581 | 0.000305176 | -1.230288045 | -6.11E-16 | 4.73E-06 | 16 |
| p_CO<sub>2</sub>_V | -4.919031267 | 0.000185345 | 0.000305176 | -1.229757817 | 4.44E-16 | 0.000654548 | 16 |
| pO<sub>2</sub>_cap | 4.888665504 | 0.000196646 | 0.000305176 | 1.222166376 | 3.55E-15 | 0.007956291 | 16 |
| CMRO<sub>2</sub> | 4.602911283 | 0.000344985 | 0.000305176 | 1.150727821 | -9.44E-16 | 0.012576333 | 16 |
| DeltaHCO<sub>2</sub> | 4.602911283 | 0.000344985 | 0.000305176 | 1.150727821 | 1.39E-16 | 0.007246483 | 16 |
| DeltaLAC | 4.602911283 | 0.000344985 | 0.000305176 | 1.150727821 | -1.33E-15 | 0.012995544 | 16 |

The metabolic concentrations are plotted in Figure 2, with and without the pseudorandom generator.

**Figure 2:**
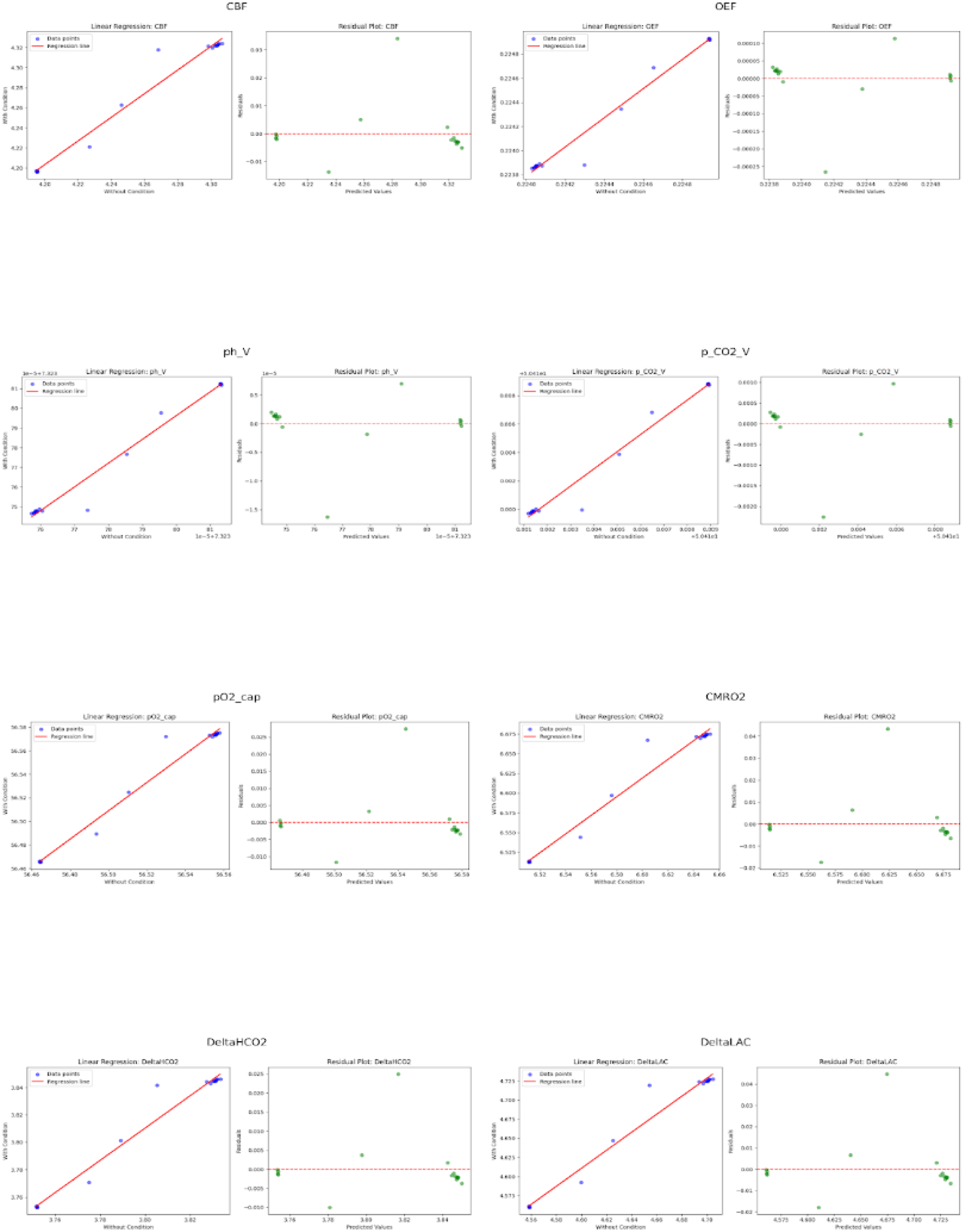
Trial 1 Metabolic Outcomes.

#### Trial 2 Outcomes

The Trial 2 prompt and response are shown in Table 3.

**Table 3:** Trial 2 Prompt and Response.

|  |  |
| --- | --- |
| Prompt: | Describe your favorite type of cuisine, including specific dishes, cultural significance, and any memorable experiences you've had with it. How does it reflect your personal tastes or background? |
| Sentence: | “Vietnamese cuisine resonates deeply with me — its balance of flavors, emphasis on fresh ingredients. Pho and banh mi stand out, embodying simplicity and depth. A meal in Hanoi, streetside, was transformative. It mirrors my preference for complexity within simplicity, a reflection of my own journey through diverse cultures and the pursuit of nuanced understanding.” |

The statistical outcomes for Trial 2 are shown in Table 4. Differences were significant (*p* < 0.001).

**Table 4:**
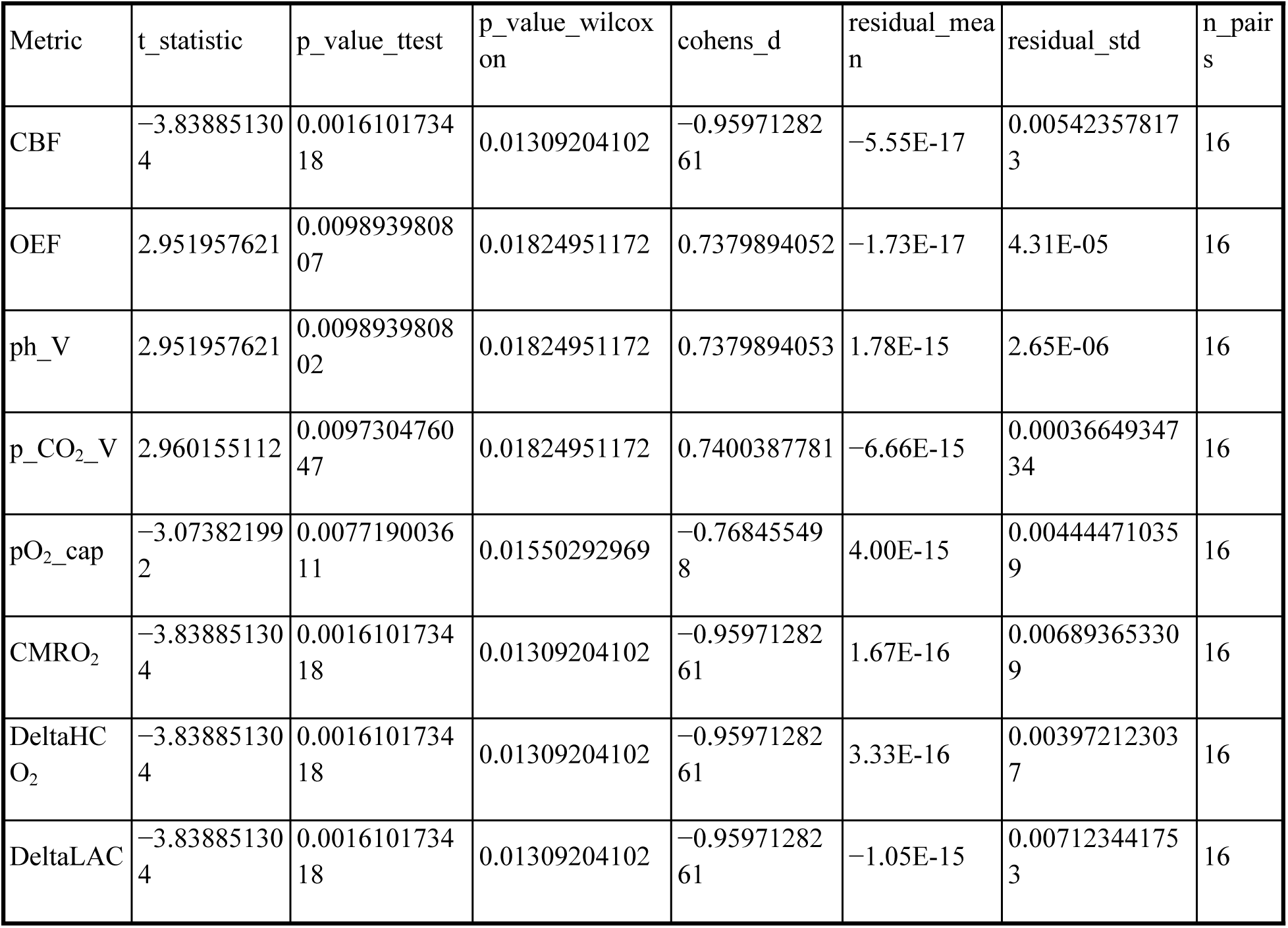
Trial 2 Statistical Outcomes.

The metabolic concentrations for Trial 2 are plotted in Figure 3, with and without the pseudorandom generator.

**Figure 3:**
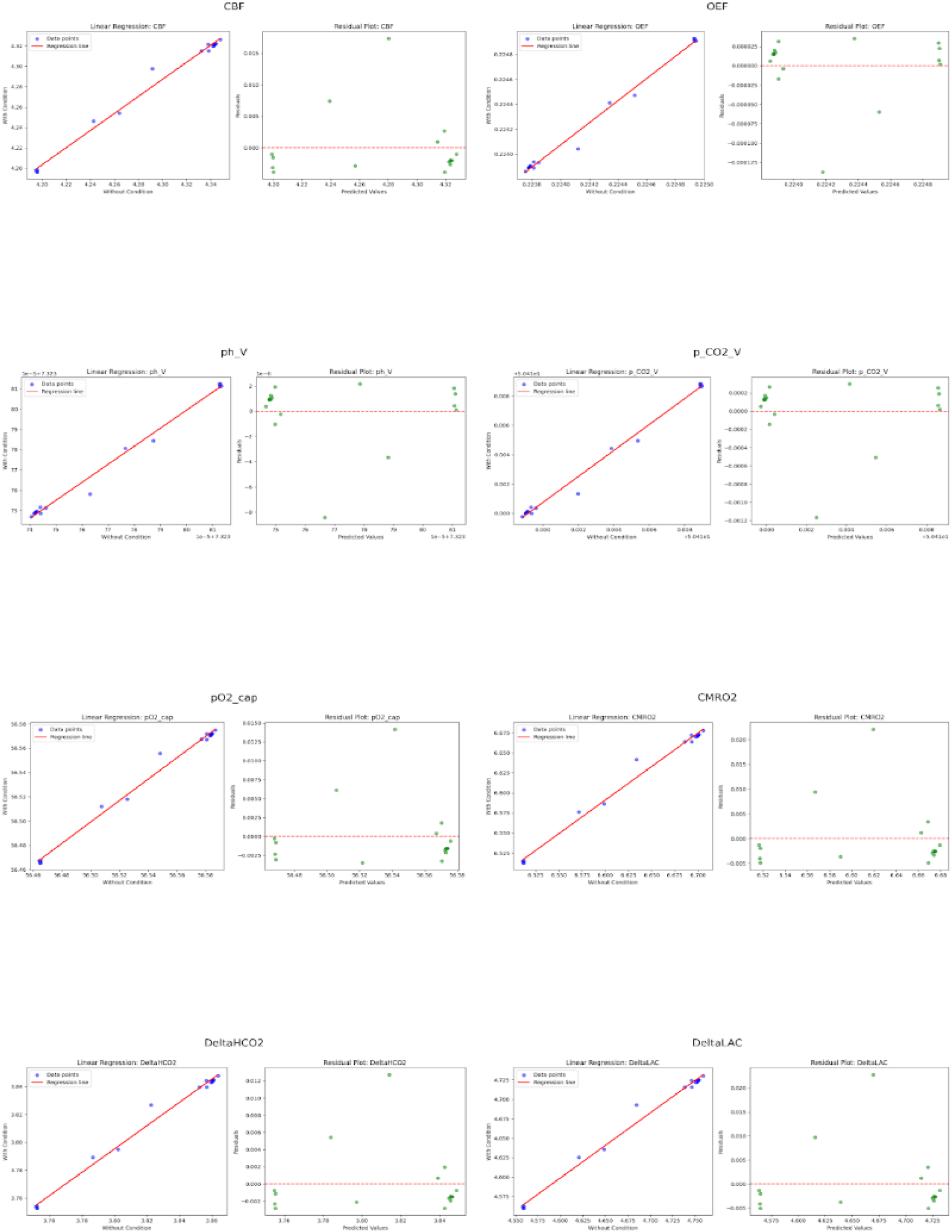
Trial 2 Metabolic Outcomes.

#### Trial 3 Outcomes

The Trial 3 prompt and response are shown in Table 5.

**Table 5:** Trial 3 Prompt and Response.

|  |  |
| --- | --- |
| Prompt: | Hello, how are you? |
| Sentence: | “Navigating through a maze of new protocols and faces, all while grappling with the humidity outside. It was invigorating yet daunting.” |

The statistical outcomes for Trial 3 are shown in Table 6. Differences were significant (*p* < 0.001).

**Table 6:** Trial 3 Statistical Outcomes.

| Metric | t_statistic | p_value_ttest | p_value_wilcoxon | cohens_d | residual_mean | residual_std | n_pairs |
| --- | --- | --- | --- | --- | --- | --- | --- |
| CBF | 7.083505189 | 3.72E-06 | 3.05E-05 | 1.770876297 | 2.22E-16 | 0.01306697457 | 16 |
| OEF | -7.232463653 | 2.91E-06 | 3.05E-05 | -1.808115913 | 1.39E-17 | 0.0001097960766 | 16 |
| ph_V | -7.232463653 | 2.91E-06 | 3.05E-05 | -1.808115913 | -2.22E-16 | 6.73E-06 | 16 |
| p_CO<sub>2</sub>_V | -7.231436863 | 2.92E-06 | 3.05E-05 | -1.807859216 | 2.66E-15 | 0.0009323334525 | 16 |
| pO<sub>2</sub>_cap | 7.216808303 | 2.99E-06 | 3.05E-05 | 1.804202076 | -4.44E-15 | 0.01126294115 | 16 |
| CMRO<sub>2</sub> | 7.083505189 | 3.72E-06 | 3.05E-05 | 1.770876297 | 5.00E-16 | 0.01660881241 | 16 |
| DeltaHCO<sub>2</sub> | 7.083505189 | 3.72E-06 | 3.05E-05 | 1.770876297 | -1.11E-16 | 0.009569997711 | 16 |
| DeltaLAC | 7.083505189 | 3.72E-06 | 3.05E-05 | 1.770876297 | 5.55E-16 | 0.01716243949 | 16 |

The metabolic concentrations for Trial 3 are plotted in Figure 4, with and without the pseudorandom generator.

**Figure 4:**
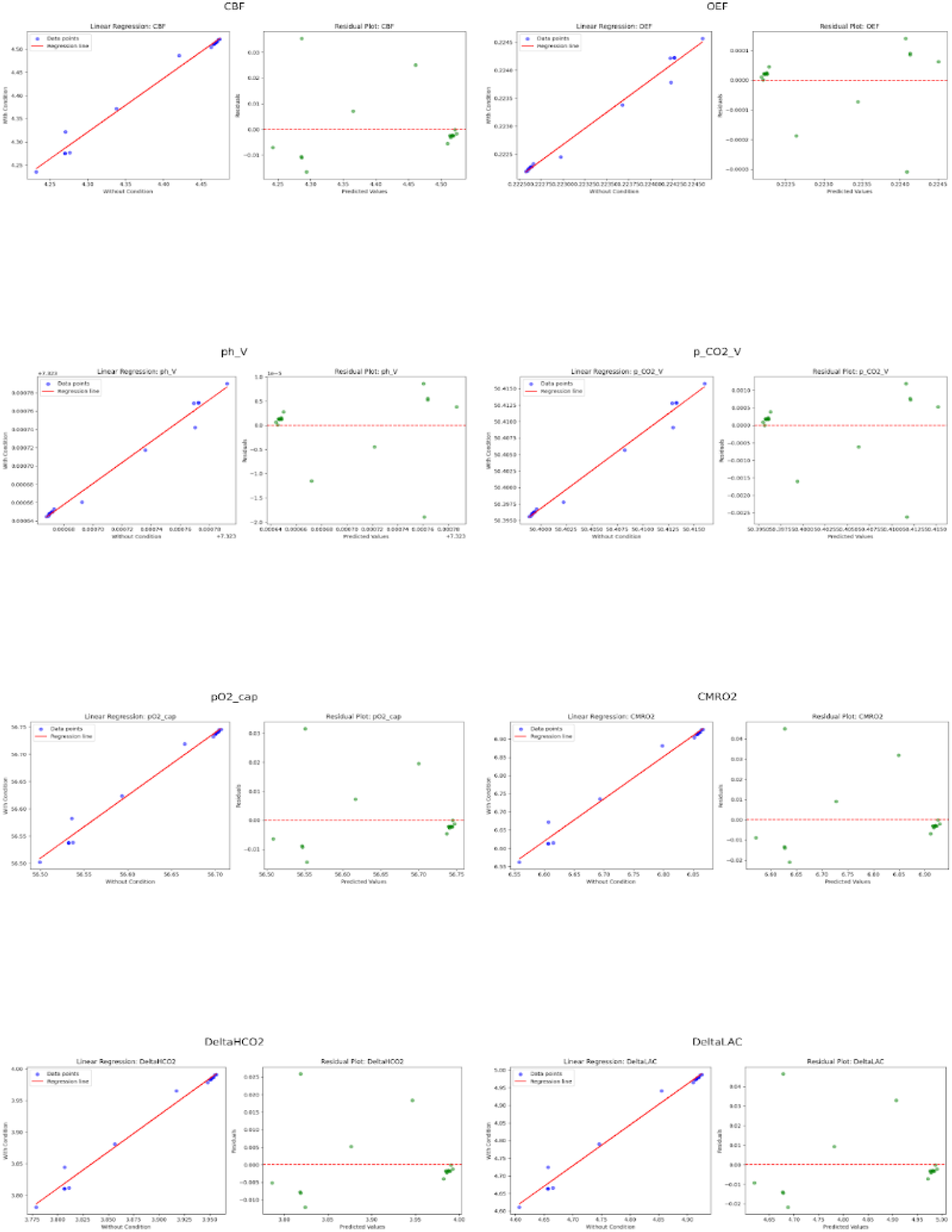
Trial 3 Metabolic Outcomes.

### GUI Panels

The GUI consisted of two primary panels:

- Text Input and Response Display: Users interacted with the GPT agent in a conversational manner. Message history was kept, which allowed for multi-turn dialogue and contextual continuance.
- Phoneme Decomposition and Visualization Control: The ARPAbet phoneme sequence generated from the GPT response was displayed for user review. From this panel, both neuroelectric and neurovascular cortical activity maps were displayed to users, who also had access to saved .mp4 versions of the visuals.

Figures 5 and 6 present screenshots of the interface at various stages of the pipeline, demonstrating the level of interaction that was provided to users.

**Figure 5:**
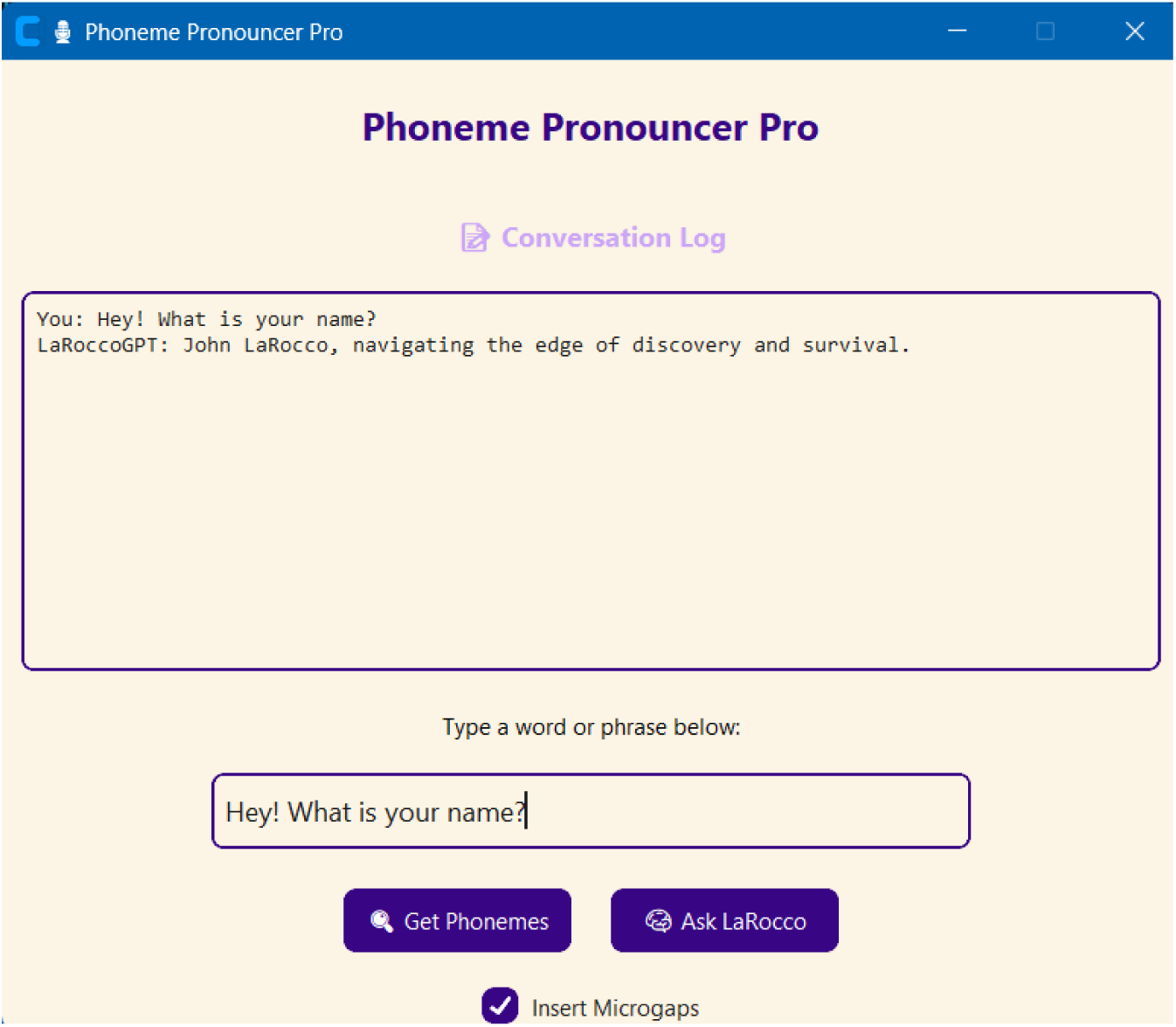
Text Input and Response Display GUI Screenshot.

**Figure 6:**
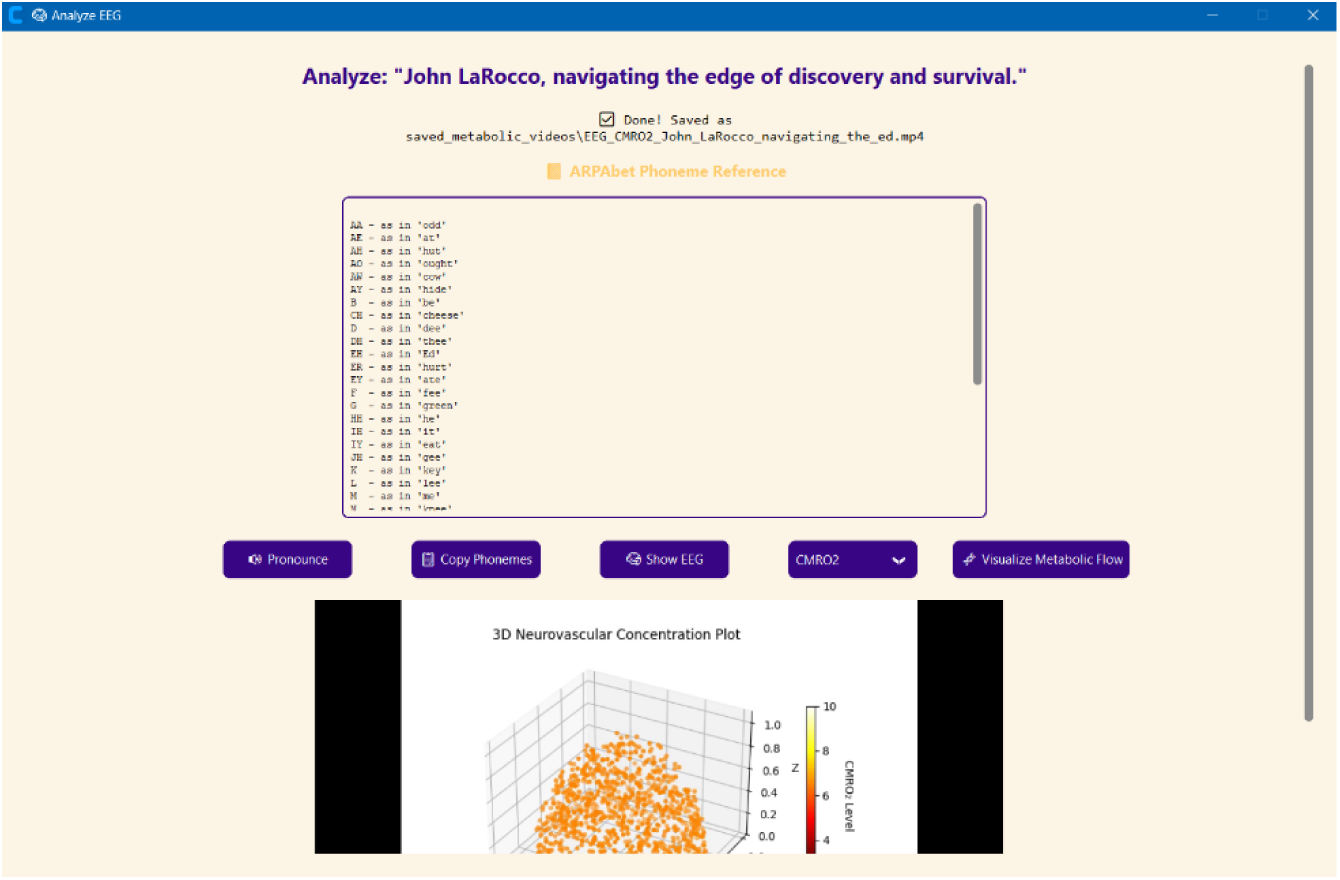
Phoneme Decomposition and Visualization Control GUI Screenshot.

### Brain Visualization

Visualization outputs were generated from each trial to demonstrate the dynamic spatial and temporal responses from the pipeline. Each input resulted in metabolic cortical activity, which was reconstructed frame-by-frame based on the synthesized EEG signals, using both electric and vascular modeling. Electrical activity is shown in Figure 7, PO_2_ modeling is shown in Figure 8, and HCO_2_ modeling is shown in Figure 9.

**Figure 7:**
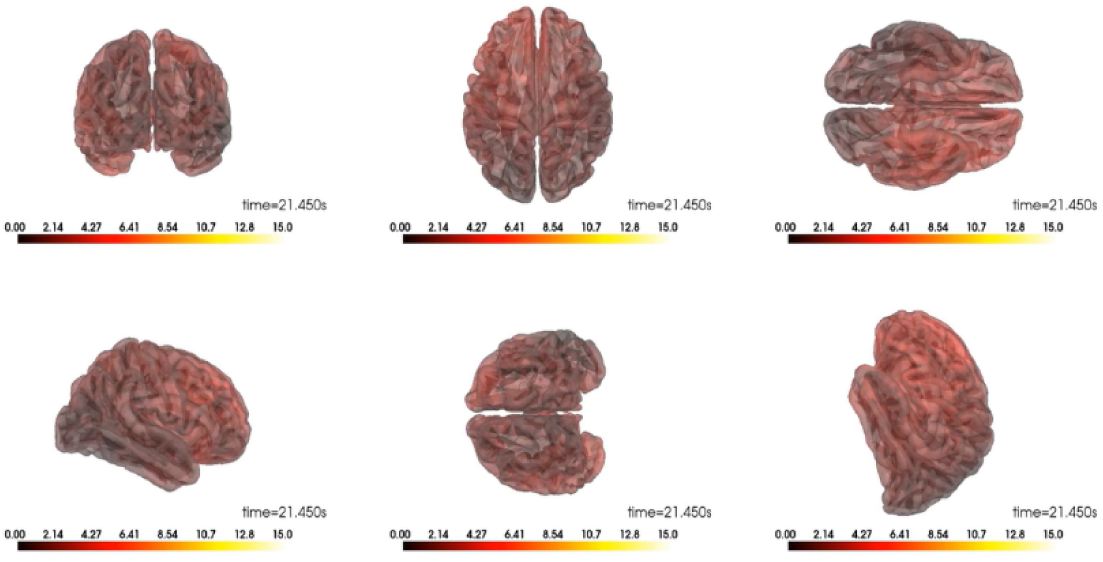
MNE Electrical Activity Plots.

**Figure 8:**
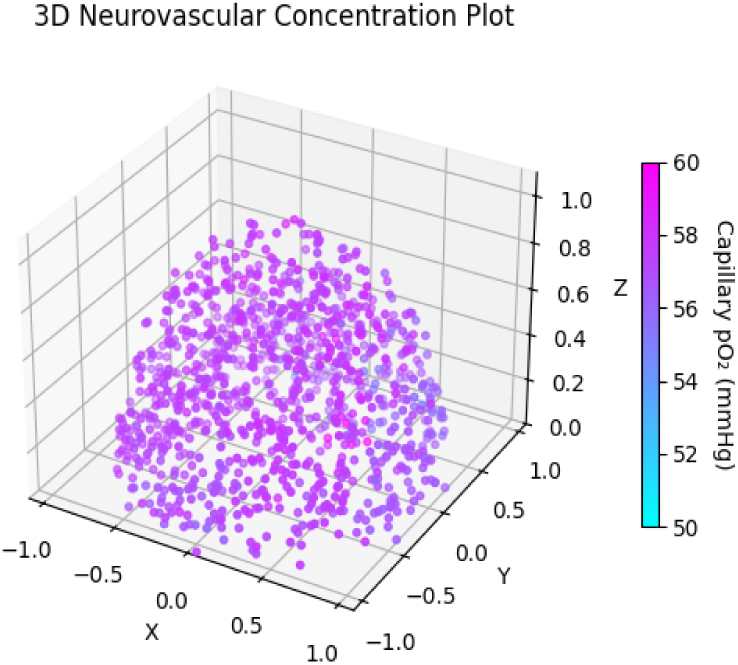
Modeling PO_2__cap in 3D.

**Figure 9:**
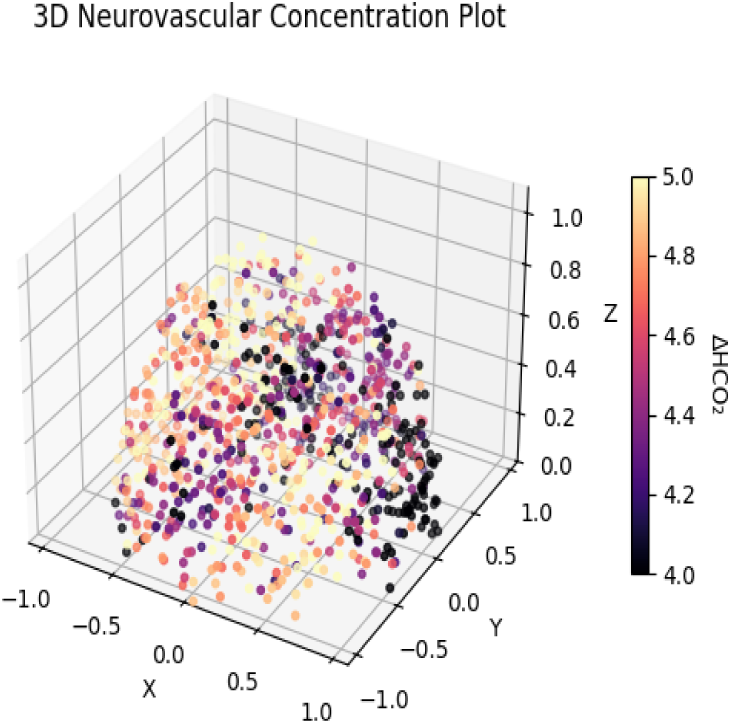
Modeling HCO_2_ in 3D.

## Discussion

### Interpretation

Including a pseudorandom generator with the existing biological models significantly altered the results (*p* < 0.001). Prior work in EEG-based ISR lacked the integration of LLMs and a broad range of biologically inspired variability (Pliatsikas et al., 2021; Guan, 2024; Tang et al., 2024). The added variability was deterministic, based entirely on pre-recorded EEG data. The ‘fractal’ nature of the simulation is also important, allowing the resolution of the visualization and parameters to be scaled depending on the available computing power, from mobile devices to supercomputers. Furthermore, the system’s modularity enables any, or all, components to be changed for different requirements. Simulating the metabolic costs of neurolinguistic processing is one application, but speech-related pathologies and the cognitive process can be simulated with minor adjustments (Pliatsikas et al., 2021; Tang et al., 2024). Ultimately, the integration of an LLM and biological variance can be used to simulate the process of sentence formation, from higher levels of abstraction (like ISR) down to molecular-level insights.

### Limitations

The current model possesses several limitations. First, a single LLM model was used for most of the processing, which still requires online access. Likewise, running a GPT model is a hardware-intensive process, but is not essential to the system. Second, using data based on a single individual from a healthy controls study, rather than a population, limited the potential for investigating speech or neural pathologies (LaRocco et al., 2023). Third, the variability of the model was pseudorandom instead of truly random, but this was only necessary to work around the constraints of the prior dataset (LaRocco et al., 2023). Finally, the software was largely based on Python, and it was not optimized for all hardware; however, using Python enabled a direct and rapid deployment of the system. Despite these limitations, the proposed system provides a template for future development.

### Future Work

The proposed ISR system requires further development. The GPT model can be adapted to run locally on less computational resources, or omitted entirely. A large population of individuals, including those with different writing styles and languages, can be used for modeling a wider range of linguistic capabilities. Adding emotional context and syntax to event responses may also improve naturalistic responses. A broader range of physiological data can be integrated, including both healthy individuals and those with neuropathologies, augmented with genuinely random data. Other types of neuroimaging data can also be integrated. The software can be optimized substantially, so it can run on a broader range of devices and networks, including edge computing. The system can potentially be extended to hardware with neural phantoms (LaRocco et al., 2024). The system can also be adapted for various hardware applications like human-computer interaction, bioreactor management, and AI agents.

## Conclusions

Despite certain limitations—such as the reliance on a single LLM, dependence on online access, use of data from a single individual, and hardware constraints—the presented system establishes a robust template for future research and development. This study demonstrated that integrating a pseudorandom generator and LLM into biologically inspired EEG-based ISR systems produces significantly altered and more flexible results compared with previous approaches. By introducing scalable, fractal-like variability and modular components, the system supports a wide range of applications—from simulating neurolinguistic metabolic costs to modeling speech pathologies and cognitive processes. The ability to adjust the resolution and parameters according to available computational resources further enhances the system’s versatility, making it suitable for deployment on devices ranging from mobile phones to supercomputers. Looking forward, the ISR system can be expanded by incorporating locally run or lighter-weight language models, integrating data from more diverse populations, and employing genuinely random variability. Optimizing the software for broader hardware compatibility and exploring hardware implementations, such as neural phantoms, emotional context, or human-computer interaction platforms, represent promising avenues for further development. Overall, this work lays the groundwork for a new generation of biologically inspired, modular, and highly adaptable ISR systems for research and practical applications.

## Acknowledgments

The authors would like to thank Prof. David Tomasko and Ron Vlcek from Ohio State University.

## Data Availability

The data and software can be found at: https://github.com/neurotechclubosu/thinktank-clearmind.

## Conflict of Interest Statement

The authors have no conflicts of interest to declare.

## Ethics Statement

No live subjects were used in this study.

## Notes

### Competing Interest Statement

The authors have declared no competing interest.

## References

Ahmadova, S. (2024) ‘Mental spaces: The cognitive structure of human thought’, Filologiya m., (3), pp. 245. 10.62837/2024.3.245

Anderson, A. et al. (2021) ‘Deep artificial neural networks reveal a distributed cortical network encoding propositional sentence-level meaning’, Journal of Neuroscience, 41(18), pp. 4100–4119. 10.1523/jneurosci.1152-20.2021

ARPAbet: A phonetic transcription system used to represent the sounds of spoken English (American Psychological Association, n.d.).

Aubinet, C. et al. (2020) ‘Brain metabolism but not gray matter volume underlies the presence of language function in the minimally conscious state (MCS): MCS+ versus MCS− neuroimaging differences’, Neurorehabilitation and Neural Repair, 34(2), pp. 172–184. 10.1177/1545968319899914

Aubinet, C. et al. (2019) ‘Reappearance of command-following is associated with the recovery of language and internal-awareness networks: A longitudinal multiple-case report’, Frontiers in Systems Neuroscience, 13. 10.3389/fnsys.2019.00008

Burgaleta, M. et al. (2016) ‘Bilingualism at the core of the brain: Structural differences between bilinguals and monolinguals revealed by subcortical shape analysis’, Neuroimage, 125, pp. 437–445. 10.1016/j.neuroimage.2015.09.073

Buxton, R. B. (2024) ‘Thermodynamic limitations on brain oxygen metabolism: Physiological implications’, The Journal of Physiology, 602(4), pp. 683–712.

Cahana-Amitay, D. et al. (2012) ‘Effects of hypertension and diabetes on sentence comprehension in aging’, The Journals of Gerontology Series B, 68(4), pp. 513–521. 10.1093/geronb/gbs085

Cahana-Amitay, D. et al. (2015) ‘Effects of metabolic syndrome on language functions in aging’, Journal of the International Neuropsychological Society, 21(2), pp. 116–125. 10.1017/s1355617715000028

Camandola, S. and Mattson, M. (2017) ‘Brain metabolism in health, aging, and neurodegeneration’, The Embo Journal, 36(11), pp. 1474–1492. 10.15252/embj.201695810

Caucheteux, C., Gramfort, A. and King, J. (2023) ‘Evidence of a predictive coding hierarchy in the human brain listening to speech’, Nature Human Behaviour, 7(3), pp. 430–441. 10.1038/s41562-022-01516-2

Chubaryan, A. and Vardanyan, M. (2024) ‘Exploring the cognitive dimensions of language acquisition’, Armenian Folia Anglistika, 20(1 (29)), pp. 13–24. 10.46991/afa/2024.20.1.13

Clercq, P. et al. (2024) ‘Neural substrates and behavioral relevance of speech envelope tracking: Evidence from post-stroke aphasia’. 10.1101/2024.03.26.586859

Cogitate C. et al. (2025) ‘Adversarial testing of global neuronal workspace and integrated information theories of consciousness’, Nature, pp. 1–10.

Crouzet, C. et al. (2016) ‘Cerebral blood flow is decoupled from blood pressure and linked to EEG bursting after resuscitation from cardiac arrest’, Biomedical Optics Express, 7(11), pp. 4660. 10.1364/boe.7.004660

Devauchelle, A. et al. (2009) ‘Sentence syntax and content in the human temporal lobe: An fMRI adaptation study in auditory and visual modalities’, Journal of Cognitive Neuroscience, 21(5), pp. 1000–1012. 10.1162/jocn.2009.21070

Devlin, J., Matthews, P. and Rushworth, M. (2003) ‘Semantic processing in the left inferior prefrontal cortex: A combined functional magnetic resonance imaging and transcranial magnetic stimulation study’, Journal of Cognitive Neuroscience, 15(1), pp. 71–84. 10.1162/089892903321107837

Ding, J. et al. (2021) ‘A metabolome atlas of the aging mouse brain’, Nature Communications, 12(1). 10.1038/s41467-021-26310-y

DiNuzzo, M. et al. (2024) ‘Neurovascular coupling is optimized to compensate for the increase in proton production from nonoxidative glycolysis and glycogenolysis during brain activation and maintain homeostasis of pH, p CO_2_, and p O_2_’,. Journal of Neurochemistry, 168(5), pp. 632–662.

Dong, X. et al. (2021) ‘Development and validation of auto-neo-electroencephalography (EEG) to estimate brain age and predict report conclusion for electroencephalography monitoring data in neonatal intensive care units’, Annals of Translational Medicine, 9(16), pp. 1290–1290. 10.21037/atm-21-1564

Fedorenko, E. et al. (2016) ‘Neural correlate of the construction of sentence meaning’, Proceedings of the National Academy of Sciences, 113(41). 10.1073/pnas.1612132113

Fedorenko, E. and Thompson-Schill, S. (2014) ‘Reworking the language network’, Trends in Cognitive Sciences, 18(3), pp. 120–126. 10.1016/j.tics.2013.12.006

Fedorenko, E., Behr, M. and Kanwisher, N. (2011) ‘Functional specificity for high-level linguistic processing in the human brain’, Proceedings of the National Academy of Sciences, 108(39), pp. 16428–16433. 10.1073/pnas.1112937108

Feng, W. (2024) ‘A review of embodied linguistics research’, Lecture Notes on Language and Literature, 7(4). 10.23977/langl.2024.070401

Friederici, A. (2004) ‘Event-related brain potential studies in language’, Current Neurology and Neuroscience Reports, 4(6), pp. 466–470. 10.1007/s11910-004-0070-0

Ghijsen, M. et al. (2018) ‘Quantitative real-time optical imaging of the tissue metabolic rate of oxygen consumption’, Journal of Biomedical Optics, 23(03), 1. 10.1117/1.jbo.23.3.036013

Gramfort, A. et al. (2013) ‘MEG and EEG data analysis with MNE-Python’, Frontiers in Neuroscience, 7(267), pp. 1–13. 10.3389/fnins.2013.00267

Goddard, C. (2020) Cognitive linguistics, pp. 1–5. 10.1002/9781118786093.iela0059

Goyal, M. et al. (2023) ‘Brain aerobic glycolysis and resilience in alzheimer disease’, Proceedings of the National Academy of Sciences, 120(7). 10.1073/pnas.2212256120

Guan, S. (2024) ‘Research on the reconstruction of language teaching cognitive process by the advanced achievements in neurolinguistics’, Journal of Educational Research and Policies, 6(7), pp. 94–99. 10.53469/jerp.2024.06(07).20

Heuven, W., Schriefers, H., Dijkstra, T. and Hagoort, P. (2008) ‘Language conflict in the bilingual brain’, Cerebral Cortex, 18(11), pp. 2706–2716. 10.1093/cercor/bhn030

Holubenko, N. and Demetskaya, V. (2020) ‘Category of modality through the prism of multipole approaches in the modern translation theory’, Journal of History Culture and Art Research, 9(2), pp. 303. 10.7596/taksad.v9i2.2500

Hyder, F. and Rothman, D. (2017) ‘Advances in imaging brain metabolism’, Annual Review of Biomedical Engineering, 19(1), pp. 485–515. 10.1146/annurev-bioeng-071516-044450

Imaezue, G. (2017) ‘Brain localization and the integrated systems hypothesis: Evidence from Broca’s region’, Journal of Behavioral and Brain Science, 07(11), pp. 511–519. 10.4236/jbbs.2017.711036

Jacob, A. et al. (2019) ‘Extracellular cold inducible RNA-binding protein mediates binge alcohol-induced brain hypoactivity and impaired cognition in mice’, Molecular Medicine, 25(1). 10.1186/s10020-019-0092-3

Kleinridders, A. et al. (2018) ‘Regional differences in brain glucose metabolism determined by imaging mass spectrometry’, Molecular Metabolism, 12, pp. 113–121. 10.1016/j.molmet.2018.03.013

Képes, Z. et al. (2021) ‘Glucose-level dependent brain hypometabolism in type 2 diabetes mellitus and obesity’, European Journal of Hybrid Imaging, 5(1). 10.1186/s41824-021-00097-z

Koush, Y. et al. (2022) ‘Human brain functional MRS reveals interplay of metabolites implicated in neurotransmission and neuroenergetics’, Journal of Cerebral Blood Flow & Metabolism, 42(6), pp. 911–934.

Krishnan, A. and Gandour, J. (2009) ‘The role of the auditory brainstem in processing linguistically-relevant pitch patterns’, Brain and Language, 110(3), pp.135–148. 10.1016/j.bandl.2009.03.005

Kuntso, O. (2022) ‘PROFILING IN LITERARY TEXT’, Collection of scientific papers «ΛΌГOΣ», (May 20, 2022; Cambridge, United Kingdom), pp. 208–209. 10.36074/logos-20.05.2022.063

LaRocco, J. et al. (2023) ‘Evaluation of an English language phoneme-based imagined speech brain computer interface with low-cost electroencephalography’, Frontiers in neuroinformatics, 17, pp. 1306277.

LaRocco, J. et al. (2024) ‘Evaluating commercial electrical neuromodulation devices with low-cost neural phantoms’, Applied Sciences, 14(14), pp. 6328.

Le, M. and Nguyen, T. (2023) ‘Roles of cognitive linguistics to second language acquisition’, Icte Conference Proceedings, 3, pp. 118–126. 10.54855/ictep.2339

Lee, S. et al. (2019) ‘Quantitative EEG predicts outcomes in children after cardiac arrest’, Neurology, 92(20). 10.1212/wnl.0000000000007504

Li, X. (2025) ‘Quantitative mapping of key glucose metabolic rates in the human brain using dynamic deuterium magnetic resonance spectroscopic imaging’, Pnas Nexus, 4(3). 10.1093/pnasnexus/pgaf072

Li, X. et al. (2020) ‘Positron emission tomography/computed tomography dual imaging using 18-fluorine flurodeoxyglucose and 11c-labeled 2-β-carbomethoxy-3-β-(4-fluorophenyl) tropane for the severity assessment of Parkinson disease’, Medicine, 99(14), e19662. 10.1097/md.0000000000019662

Lin, Q. (2024) ‘Exploration of conceptual metaphor theory in cognitive linguistics’, Lecture Notes on Language and Literature, 7(1). 10.23977/langl.2024.070113

Loh, W. L. (1996) ‘On Latin hypercube sampling’, The annals of statistics, 24(5), pp. 2058–2080.

Mao, H. et al. (2024) ‘1h-magnetic resonance spectroscopy (MRS) in analysis of cerebral metabolite: MRS image for elderly patient anesthesia’. 10.21203/rs.3.rs-5406479/v1

Mason, R., Just, M., Keller, T. and Carpenter, P. (2003) ‘Ambiguity in the brain: what brain imaging reveals about the processing of syntactically ambiguous sentences’, Journal of Experimental Psychology Learning Memory and Cognition, 29(6), pp. 1319–1338. 10.1037/0278-7393.29.6.1319

Matthews, D. et al. (2018) ‘FDG PET Parkinson’s disease-related pattern as a biomarker for clinical trials in early stage disease’, Neuroimage Clinical, 20, pp. 572–579. 10.1016/j.nicl.2018.08.006

McConnell, B. et al. (2024) ‘On monitoring brain health from the depths of sleep: Feature engineering and machine learning insights for digital biomarker development’. 10.1101/2024.02.27.581950

Müller, M. et al. (2020) ‘Standardized visual EEG features predict outcome in patients with acute consciousness impairment of various etiologies’, Critical Care, 24(1). 10.1186/s13054-020-03407-2

Nelson, M. et al. (2017) ‘Neurophysiological dynamics of phrase-structure building during sentence processing’, Proceedings of the National Academy of Sciences, 114(18). 10.1073/pnas.1701590114

OpenAI. (2025). *ChatGPT* (Mar 14 version) [Large language model]. https://chat.openai.com/chat

Pang, H. et al. (2020) ‘Precision mapping of the mouse brain metabolome’. 10.1101/2020.12.28.424544

Pinti, P.. (2021) ‘An analysis framework for the integration of broadband NIRS and EEG to assess neurovascular and neurometabolic coupling’, Scientific Reports, 11(1). 10.1038/s41598-021-83420-9

Pliatsikas, C. and Luk, G. (2016) ‘Executive control in bilinguals: A concise review on fmri studies’, Bilingualism Language and Cognition, 19(4), pp. 699–705. 10.1017/s1366728916000249

Pliatsikas, C. et al. (2021) ‘Bilingualism is a long-term cognitively challenging experience that modulates metabolite concentrations in the healthy brain’, Scientific Reports, 11(1). 10.1038/s41598-021-86443-4

Price, C. (2012) ‘A review and synthesis of the first 20 years of pet and fmri studies of heard speech, spoken language and reading’, Neuroimage, 62(2), pp. 816–847. 10.1016/j.neuroimage.2012.04.062

Pylkkänen, L. and McElree, B. (2007) ‘An MEG study of silent meaning’, Journal of Cognitive Neuroscience, 19(11), pp. 1905–1921. 10.1162/jocn.2007.19.11.1905

Qin, J., Liu, D. and Lei, L. (2024) ‘Cognitive linguistics-inspired language instruction’, Language Teaching, 57(4), pp. 478–500. 10.1017/s0261444824000119

Radder, N., Sonar, S., Nanivadekar, A. and Radder, S. (2024) ‘Synergy in neuroimaging: PET-CT and MRI fusion for enhanced characterization of brain pathology’, Cureus. 10.7759/cureus.74353

Rae, C.D. et al. (2024) ‘Brain energy metabolism: A roadmap for future research’, Journal of Neurochemistry, 168(5), pp. 910–954.

Rothman, D. L. et al. (2021) ‘Methods| 13C MRS measurements of in vivo rates of the glutamate/glutamine and GABA/glutamine neurotransmitter cycles’.

Sala, A. et al. (2021) ‘Lifelong bilingualism and mechanisms of neuroprotection in alzheimer dementia’, Human Brain Mapping, 43(2), pp. 581–592. 10.1002/hbm.25605

Saveliev, S. and Chuev, A. (2023) ‘Difficulty of energy expenditure of brain when using written speech’.

Schamberg, G., Chakravarty, S., Baum, T. and Brown, E. (2020) ‘Inferring neural dynamics during burst suppression using a neurophysiology-inspired switching state-space model’, pp. 92–96. 10.1109/ieeeconf51394.2020.9443373

Schuhmann, T., Schiller, N., Goebel, R. and Sack, A. (2009) ‘The temporal characteristics of functional activation in Broca’s area during overt picture naming’, Cortex, 45(9), pp. 1111–1116. 10.1016/j.cortex.2008.10.013

Sebastian, R., Laird, A. and Kıran, S. (2011) ‘Meta-analysis of the neural representation of first language and second language’, Applied Psycholinguistics, 32(4), pp. 799–819. 10.1017/s0142716411000075

Shtok, N. (2021) ‘Cognitive linguistics – a historical context’, Białostockie Archiwum Językowe, (21), pp. 123–138. 10.15290/baj.2021.21.08

Tan, H., Li, X., Wei, K. and Guan, Y. (2018) ‘Study on brain glucose metabolic networks in Parkinson’s disease patients with visual spatial dysfunction by 18f-FDG PET imaging’, Traditional Medicine and Modern Medicine, 01(01), pp. 27–31. 10.1142/s2575900018500015

Tang, J. et al. (2024) ‘Imagined speech reconstruction from neural signals–An overview of sources and methods. IEEE Transactions on Instrumentation and Measurement.

Tolomeo, D. et al. (2018) ‘Chemical exchange saturation transfer mri shows low cerebral 2-deoxy-d-glucose uptake in a model of Alzheimer’s disease’, Scientific Reports, 8(1). 10.1038/s41598-018-27839-7

Tripathi, E. (2024) ‘Functional brain connectivity differences between aphasic and neurotypical brains’, International Journal on Recent and Innovation Trends in Computing and Communication, 11(9), pp. 4714–4718. 10.17762/ijritcc.v11i9.10022

Wang, C. et al. (2024) ‘Brainscale: Enabling scalable online learning in spiking neural networks’. 10.1101/2024.09.24.614728

Westover, M. et al. (2015) ‘Real-time segmentation and tracking of brain metabolic state in ICU EEG recordings of burst suppression’, pp. 330–344. 10.1017/cbo9781139941433.015

Wilson, R. et al. (2019) ‘High-speed quantitative optical imaging of absolute metabolism in the rat cortex’. 10.1101/786244

Yu, J. et al. (2023) ‘Construction of an individualized brain metabolic network in patients with advanced non-small cell lung cancer by the Kullback-Leibler divergence-based similarity method: A study based on 18f-fluorodeoxyglucose positron emission tomography’, Frontiers in Oncology, 13. 10.3389/fonc.2023.1098748

Zhou, L., Li, X. and Su, B. (2022) ‘Spatial regulation control of oxygen metabolic consumption in mouse brain’, Advanced Science, 9(34). 10.1002/advs.202204468

